# Expression variation analysis for tumor heterogeneity in single-cell RNA-sequencing data

**DOI:** 10.1101/479287

**Authors:** Emily F. Davis-Marcisak, Pranay Orugunta, Genevieve Stein-O’Brien, Sidharth V. Puram, Evanthia Roussos Torres, Alexander Hopkins, Elizabeth M. Jaffee, Alexander V. Favorov, Bahman Afsari, Loyal A. Goff, Elana J. Fertig

**Affiliations:** McKusick-Nathans Institute for Genomic Medicine, Johns Hopkins School of Medicine, Baltimore, MD, USA; Department of Oncology, Sidney Kimmel Comprehensive Cancer Center, Johns Hopkins School of Medicine, Baltimore, MD, USA; Solomon H. Snyder Department of Neuroscience, Johns Hopkins School of Medicine, Baltimore, MD, USA; Department of Otolaryngology-Head and Neck Surgery, Washington University School of Medicine, St. Louis, MO, USA; Department of Genetics, Washington University School of Medicine, St. Louis, MO, USA; Michigan Center for Translational Pathology, University of Michigan, Ann Arbor, MI, USA; Laboratory of Systems Biology and Computational Genetics, Vavilov Institute of General Genetics, Russian Academy of Sciences, Moscow, Russia; Department of Applied Mathematics and Statistics, Johns Hopkins University Whiting School of Engineering, Baltimore, MD, USA

**Keywords:** tumor heterogeneity, single-cell RNA-seq, T-cell receptor, genomics, cancer systems biology

## Abstract

Tumor heterogeneity provides a complex challenge to cancer treatment and is a critical component of therapeutic response, disease recurrence, and patient survival. Single-cell RNA-sequencing (scRNA-seq) technologies reveal the prevalence of intra-and inter-tumor heterogeneity. Computational techniques are essential to quantify the differences in variation of these profiles between distinct cell types, tumor subtypes, and patients to fully characterize intra-and inter-tumor molecular heterogeneity. We devised a new algorithm, Expression Variation Analysis in Single Cells (EVAsc), to perform multivariate statistical analyses of differential variation of expression in gene sets for scRNA-seq. EVAsc has high sensitivity and specificity to detect pathways with true differential heterogeneity in simulated data. We then apply EVAsc to several public domain scRNA-seq tumor datasets to quantify the landscape of tumor heterogeneity in several key applications in cancer genomics, i.e. immunogenicity, cancer subtypes, and metastasis. Immune pathway heterogeneity in hematopoietic cell populations in breast tumors corresponded to the amount diversity present in the T-cell repertoire of each individual. In head and neck squamous cell carcinoma (HNSCC) patients, we found dramatic differences in pathway dysregulation across basal primary tumors. Within the basal primary tumors we also identified increased immune dysregulation in individuals with a high proportion of fibroblasts present in the tumor microenvironment. Moreover, cells in HNSCC primary tumors had significantly more heterogeneity across pathways than cells in metastases, consistent with a model of clonal outgrowth. These results demonstrate the broad utility of EVAsc to quantify inter-and intra-tumor heterogeneity from scRNA-seq data without reliance on low dimensional visualization.

## 1. INTRODUCTION

Tumor heterogeneity poses significant challenges in the clinical diagnosis and treatment of cancer. Variation can occur among tumors of the same histological subtype, giving rise to variability in therapeutic responses among patients. Cellular heterogeneity can also occur within tumors, allowing cancer to evolve over the course of disease progression, resulting in drug resistance, treatment failure, and disease recurrence^1–3^. An important source of tumor heterogeneity is the molecular variation among subclones and even individual cells within a tumor. This variation drives tumor progression through dysregulation of key cancer pathways and contributes to the evolutionary fitness of tumors^3,4^. Differential variability analysis of bulk transcriptional data from microarrays and RNA-sequencing have also demonstrated that tumors with worse prognosis have a corresponding increase in transcriptional variation^5–8^. Single-cell RNA-sequencing (scRNA-seq) technologies provide an unprecedented ability to measure gene expression from individual cells, enabling in-depth exploration of tumor heterogeneity^9,10^.

Accurate characterization of inter-sample variation from scRNA-seq data of tumors is critical to quantify tumor heterogeneity. Molecular heterogeneity of scRNA-seq data is often analyzed visually, using computational methods for dimensionality reduction that enable qualitative interpretations based upon the dissimilarity in transcriptional profiles between cells^11–19^. These techniques enable visualization of the cellular composition within each sample as a measure of heterogeneity. However, stochastisticity, overplotting, and nonlinearity can challenge biological interpretation from visual analysis of scRNA-seq data. Moreover, the embeddings produced by some algorithms such as tSNE do not specifically preserve cluster heterogeneity. Instead, robust statistics are essential. Coefficient of variation (CV)^20^ has been broadly applied to extend this visualization at a sample level to quantify transcript variability across samples from one group. Similarly, phenotypic volume was introduced to quantify the variation between cells in a single sample^21^. These methods are able to visually segregate groups and identify highly variable genes or samples. Additional analysis techniques are essential to capture relevant pathway level heterogeneity that drives the observed deviations between groups of cells from different phenotypes.

In this paper, we extend our algorithm to quantify relative pathway dysregulation between experimental conditions from bulk transcriptional data^22^ called Expression Variation Analysis (EVA) to scRNA-seq. We call this Expression Variation Analysis in Single Cells (EVAsc). Briefly, EVAsc provides a robust statistical test to compare the heterogeneity of transcriptional profiles of genes in a pathway between groups of cells from two phenotypes. Using simulated data, we demonstrate that this method is robust for imputed scRNA-seq data. With the recent outpouring of large scale scRNA-seq studies in cancer, publicly available datasets provide a breadth of transcriptional data to explore the role of heterogeneity in a variety of contexts. We utilize datasets from head and neck^23^ and breast^21^ cancers, which contain thousands of cells comprising dozens of cell types from different tissues, subtypes, and individuals. These datasets were selected to benchmark the performance of our algorithm to characterize cases with known differences in heterogeneity, such as between tumor and normal cells. Pathways found to be statistically significant from EVAsc are called differentially variable or heterogeneous between cells from distinct sample groups. These analyses enable novel characterization of the role of tumor heterogeneity in complex processes in cancer. For example, these analyses enable us for the first time to define the relationship between variation in immune pathways and TCR clonality. They also quantify inter-tumor heterogeneity between primary tumors of a single subtype and identify immune dysregulation related to the degree of fibroblasts present in the tumor microenvironment (TME). Finally, these analyses enable quantification of pervasive, differentially variable pathways between primary tumors and metastases consistent with the hypothesis of clonal outgrowth. Together, these results suggest that EVAsc provides an important tool to quantify inter-cellular heterogeneity directly from scRNA-seq data to yield novel biological insights that are independent of more subjective visualization techniques.

## 2. METHODS

### 2.1 EVA-sc analysis

We use EVA from the R/Bioconductor package GSReg^22^ version 1.17.0 to quantify pathway dysregulation in sets of cells from one group relative to the set of cells in another. Kendall-tau dissimilarities are computed with the function in the GSReg package and other dissimilarity measures using the R package philentropy version 0.2.0. Imputed scRNA-seq data are input to this algorithm, with imputation method described for each dataset below. Analyses are performed for gene sets for Hallmark gene set pathways from MSigDB version 6.1^24^, meta-signatures from Puram et al.^23^, and Myeloid Innate Immunity Panel pathways from nanoString (NanoString Technologies). P-values obtained from EVA analysis are FDR adjusted with the Benjamini-Hochberg correction and FDR adjusted p-values below 0.05 are called statistically significant.

All code for the EVAsc analyses is available from https://github.com/edavis71/scEVA.

### 2.2 Simulated data

We generate two simulated datasets to benchmark the performance of EVAsc, with varying degrees of complexity to balance controlled testing of the algorithm with the complex properties of scRNA-seq data. For the first, we simulate count data with different amounts of missing data using the squamous cell carcinoma bulk RNAseq dataset with a binary phenotype from the R/Bioconductor package GSBenchmark version 0.112.0. We randomly replace count data with specified percentages of zeros to generate multiple datasets with varying degrees of missingness. We also generate a dataset with no signal by duplicating the count data for one phenotype. Again, we randomize zeros to determine the effect on the false positive rate in data without signal. We perform 100 iterations of all randomizations and test the performance against 35 distance measures.

While random zeros can be used to examine the general effect of missing data on dissimilarity, this does not accurately capture the nature of zeros in scRNA-seq data. To explore this, we simulate scRNA-seq data generated using the R/Bioconductor package Splatter version 1.0.3^25^. We generated two simulated datasets: one with no signal and one with known differential variation to assess the dependence of EVAsc to missing data from scRNA-seq data. For the first, count data was simulated for a single group of 100 cells and 10,000 genes using default parameters. A second group was simulated under the same conditions, with the parameter for dropout = TRUE. Merging these outputs resulted in a single dataset with a population of cells equally distributed between two groups with identical transcriptomes and varying number of random zeros in one group.

We impute the simulated dataset described above with the R package Rmagic version 1.3.0^26^. To generate a synthetic scRNA-seq count data consisting of two groups with a high degree of differential variability, we then added random noise into the expression matrix by randomizing of the count data in each cell for one group to reflect pathway heterogeneity.

### 2.3 Cancer scRNA-seq datasets

We use 45,000 immune cells from eight primary breast carcinomas with matched normal breast tissue, blood, and lymph nodes along along with 27,000 T-cells with paired single-cell RNA and single-cell TCR sequencing previously described in Azizi et al.^21^. In our study, we impute the scRNA-seq data from Puram et al.^23^ with MAGIC version 0.1.0 (Python) prior to analysis^26^. The scRNA-seq dataset from Azizi et al.^21^ was previously imputed from their study using BiSCUIT^27^.

We also use scRNA-seq datasets of 6,000 cells from 18 head and neck squamous cell carcinoma (HNSCC) patients containing five sets of matched primary tumors and lymph node metastases as previously described in Puram et al.^23^. HNSCC subtypes present in the data were called using The Cancer Genome Atlas (TCGA) classification profiles from bulk data on primary cancer cells^28^. Batch effect correction was performed using the function ComBat from R/Bioconductor package sva version 3.26.0^29^, considering each patient as a batch to isolate differences between cells from distinct HNSCC subtypes.

### 2.4 TCR repertoire analysis

TCR repertoire clonality, richness, and Morisita-Horn similarity index between samples were computed on the TCR sequencing data from Azizi et al.^21^ using the R package tcrSeqR^30^ version 1.0.6 available from https://github.com/ahopki14/tcrSeqR.

### 2.5 Differential expression and gene set enrichment analysis

Differential expression analyses were performed across all expressed genes using the Monocle R/Bioconductor package version 2.6.1^31^. In all tests, the number of genes detected in each cell was included in both the full and reduced models as a nuisance parameter. Gene set enrichment was performed on differentially expressed genes with FDR adjusted p-values below 0.05 using the wilcoxGST function from the R package LIMMA version 3.32.10^32^. The alternative hypotheses of “up” and “down” were used to determine if genes within Myeloid Innate Immunity Panel pathways were generally upregulated or downregulated, respectively.

## 3. RESULTS

### 3.1 EVAsc algorithm

EVA is a statistical algorithm designed to compare the expected dissimilarity of expression profiles between all pairs of samples from one phenotype relative to the expected dissimilarity of expression profiles between all pairs of samples from another. When applied to the set of genes in a pathway, the expected dissimilarity between all pairs of samples from one phenotype provides a measure of pathway dysregulation, which we denote as the EVA statistic. EVA tests the null hypothesis that pathway dysregulation is equal in the phenotypes using a computationally efficient approximation for p-values from U-theory statistics^22^. The resulting EVA algorithm provides a robust, non-competitive gene set measure to quantify the relative inter-phenotype heterogeneity of pathway usage. In our previous applications, we based our comparisons on the Kendall-tau dissimilarity measure in bulk transcriptional data. This measure was selected both because its rank-based nature reduces sensitivity to data preprocessing and models discordance between the expression of genes in a profile, which is indicative of pathway dysregulation. Bulk data lacks the resolution to quantify cellular heterogeneity because it is inherently an aggregate. EVA is poised to perform variation analysis based upon the measures of cellular heterogeneity in scRNA-seq data. If we treat each individual cell as a sample, we can adapt EVA to compare transcriptional heterogeneity scRNA-seq data between specified sets of cells (Figure 1).

**Figure 1.**
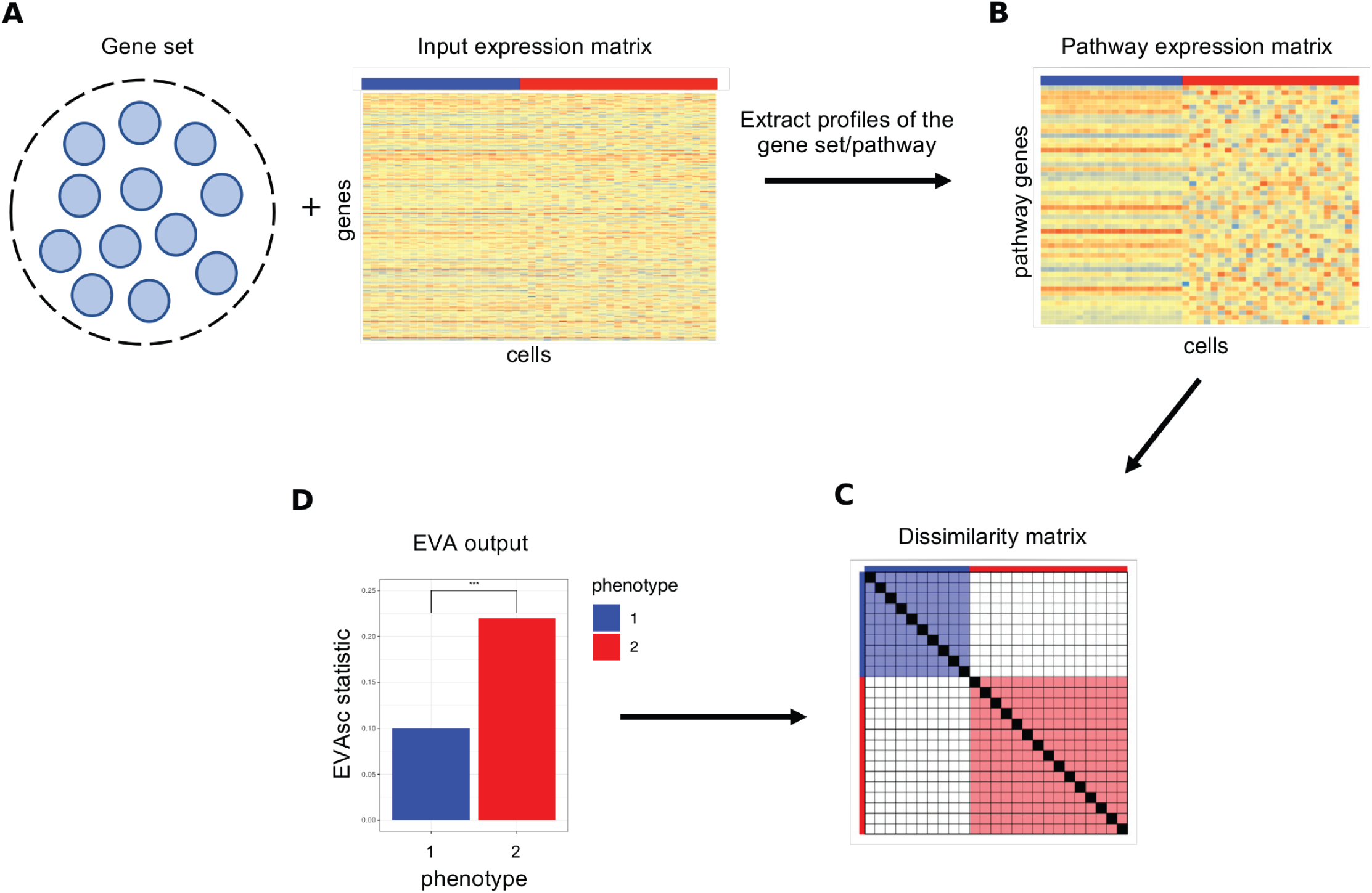
Overview of EVAsc algorithm to compare pathway-level transcriptional heterogeneity between groups of cells from two phenotypes. EVAsc inputs a single-cell gene expression matrix for cells from two phenotypes (blue and red) and a list of genes annotated to a single pathway. B. EVAsc extracts the expression profiles for pathway specific genes. C. It then computes the dissimilarity between the expression profiles for each pair of cells from the same phenotype using a user specified dissimilarity metric. D. Finally, EVAsc computes the expected dissimilarity between pairs of cells of each pheno-type and U-theory statistics are applied to test the null hypothesis that the expected dissimilarity between pairs of cells from one phenotype is equal to the expected dissimilarity between paris of cells in the other. The expected dissimilarity between pairs of cells from one phenotype is called the EVAsc statistic, which quantifies the inter-cellular heterogeneity for a given pathway. The U-theory statistics provide a robust estimate to quantify p-values that compare this relative heterogeneity between phenotypes.

Given that Kendall-tau dissimilarity is rank-based, it is robust to normalization and read depth. However, the abundance of zero counts from scRNA-seq data would lead to an increase of ties in the ranking. Moreover, dropout events in scRNA-seq data occur when an mRNA transcript is not captured by the library preparation reaction prior to sequencing and this generally happens more frequently in genes expressed at low levels. This, combined with the general bursting nature of the transcription machinery, leads to “false” zero counts, indistinguishable from biological zeros of truly unexpressed transcripts and inappropriate rank assignments in the Kendall-tau dissimilarity.

### 3.2 Simulated data

The EVAsc algorithm defaults to comparisons based upon the Kendall-tau dissimilarity metric. Because this metric quantifies the number of gene pairs which switch ranks between two conditions, it directly quantifies how tightly a set of genes in a pathway are regulated^5,22^. Yet, the U-theory statistics to compare the expected dissimilarity between groups of cells from distinct phenotypes are general and can be applied to any dissimilarity measure. In order to compare the sensitivity of different dissimilarity measures to variable sparsity, we use a bulk RNAseq dataset from GSBenchmark containing normal and tumor samples. This dataset has 50 pathways which are significantly dysregulated between tumor and normal samples in bulk. For each metric, the significant pathways calculated on the data with no sparsity are used as our true positives in the scRNA-seq simulation respectively. We then test the performance of EVAsc using 35 distance measures when varying percentages of missing data are present based upon these true positives. Even with no missing data, the number of significant pathways between tumor and normal vary widely across metrics (Supplemental Figure 1A). Several metrics including cosine and Ruzicka found no significant differentially variable pathways between normal and tumor samples. Kendall-tau detected the highest number of significant pathways, followed by Euclidean which is a commonly used distance to compare transcriptomes between single cells in methods such as tSNE^11–17^.

When the dataset is mirrored to produce two identical groups with no signal, as the percentage of zeros in the dataset increase so do the number of falsely detected significant pathways for a majority of the metrics (Supplemental Figure 1B). The number of correctly identified significant true and false pathways compared to the known ground truth in the simulated dataset with signal vary greatly depending on the amount of missing data, with an overall loss of signal when the amount of zeros is the highest (Supplemental Figure 1C and 1D). Of note, Kendall-tau resulted in the detection of the greatest number of significant pathways in data with no zeros, and the lowest false positives of all metrics with increasing zeros in datasets with or without signal. These data indicate that all metrics have varying degrees of sensitivity to missing data. We select Kendall-tau for the remainder of the analyses in this paper based on the observed accuracy in the two simulated datasets without additional normalization. We note the rank-based nature of the Kendall-tau dissimilarity renders the EVA statistics performed on Kendall-tau dissimilarity independent of common normalization procedures, such as log transformation.

To determine the effect of dropout and imputation on EVAsc’s robustness to detect pathway variability, we conducted a simulation study using synthetic scRNA-seq datasets generated using the Splatter pipeline. We first examined the performance of EVAsc on a dataset with no signal and a bias in zeros. The simulated dataset contained two identical groups, one containing only biological zeros, and one where random dropout was also present (Figure 2A). Due to the abundance of zeros in the group with dropout and the sensitivity of Kendall-tau to missing data, EVAsc failed to recognize that the groups were otherwise identical and detected differential heterogeneity across 62% (31 out of 50) MSigDB Hallmark gene set pathway comparisons. We then imputed the missing values in the simulated dataset using MAGIC^26^. EVAsc analysis of this imputed data had no pathways with statistically significant differential heterogeneity between the two groups (Figure 2B).

**Figure 2.**
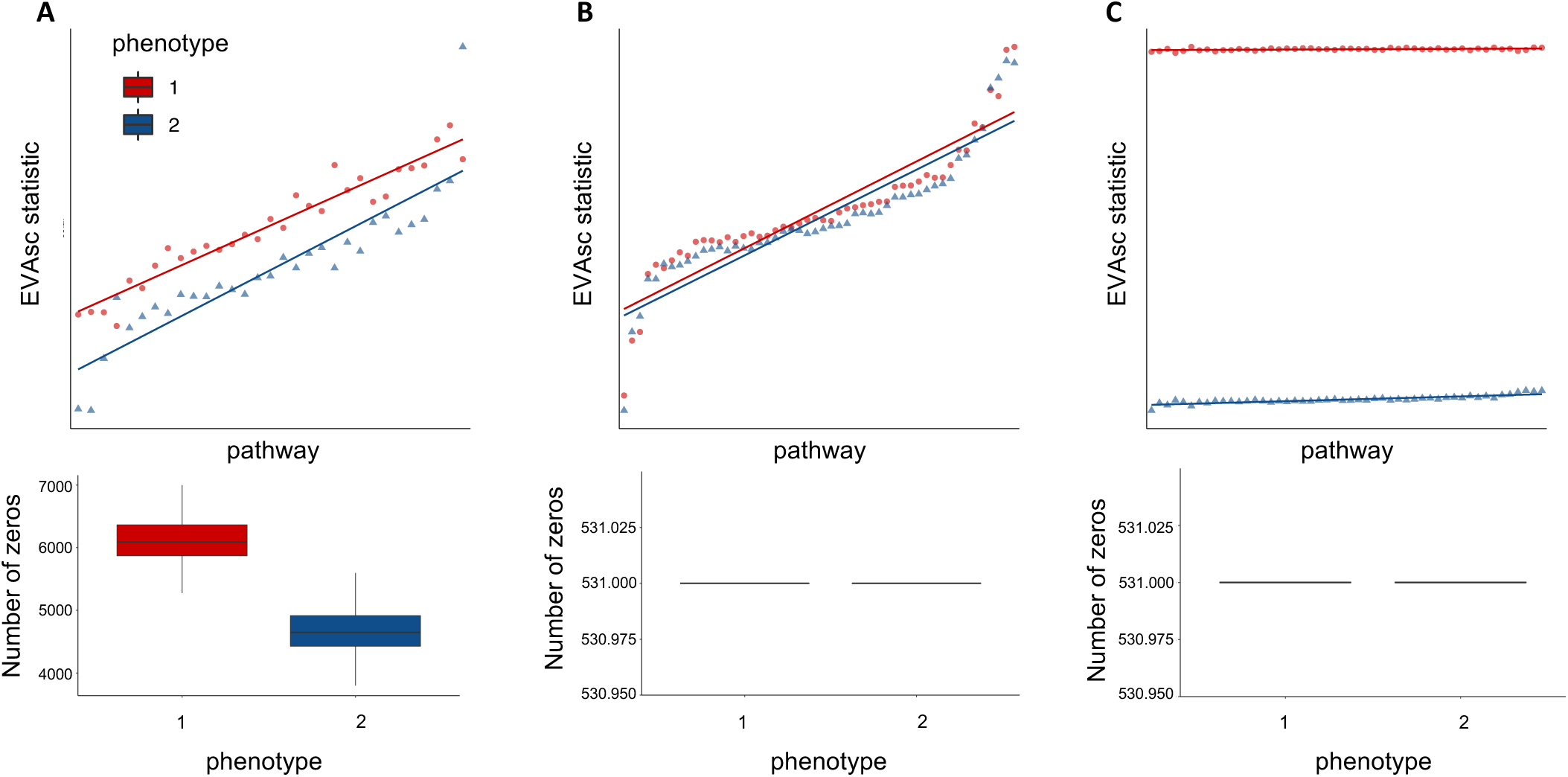
Performance of EVAsc with Kendall-tau dissimilarity on simulated data. A. We apply EVAsc to a simulated dataset containing fifty pathways with no differential variation between cells from two phenotypes, but differential bias in their respective dropout rates. EVAsc statistics using a Kend-all-tau dissimilarity have differential heterogeneity consistent with the simulated dropout rates. B. After MAGIC imputation of the data from A, EVAsc finds no significant differentially variable pathways and EVAsc statistics overlap for the two groups. C. We generate an additional simulated dataset by adding randomized signal to one group from the imputed data. The EVAsc statistics for significant pathways reflects the true heterogeneity in the simulated dataset.

We next examined the performance of EVAsc to detect known differential variation in imputed scRNA-seq data. To simulate heterogeneity, pathway expression profiles for each cell in one group were randomized from the previously described imputed dataset. EVAsc detected dramatic differences in variation between the two groups across all randomized hallmark pathways. 100% (50 out of 50) of the comparisons were statistically significant. These simulations demonstrate that EVAsc is able to assess the degree of pathway dysregulation between conditions in imputed scRNA-seq data.

### 3.3 EVAsc detects greater variation in tumor than normal in samples in a real dataset

We next evaluated the ability of EVAsc to compare heterogeneity between normal and tumor samples in scRNA-seq data from breast tumors for distinct immune cell types^21^. Azizi et al.^21^ reported an increase in the variance of tumor cell-intrinsic gene expression compared to normal breast tissue. Genes with the largest differential variance were enriched in signaling pathways important to the TME. To demonstrate that EVAsc enables robust statistical comparison of this heterogeneity in pathways, we compared tumor to normal immune cells across multiple cell types, which included T-cells, myeloid, and NK cells. EVAsc analysis detected greater variation in breast tumor than normal breast tissue across each immune cell type tested. All 50 pathways tested were statistically significant in each comparison (FDR adjusted p-value < 0.05) (Figure 3, Supplemental Tables 1-4). This suggests that increased pathway heterogeneity within tumor-associated immune cell types may be driven by distinct TMEs present within a single tumor.

**Figure 3.**
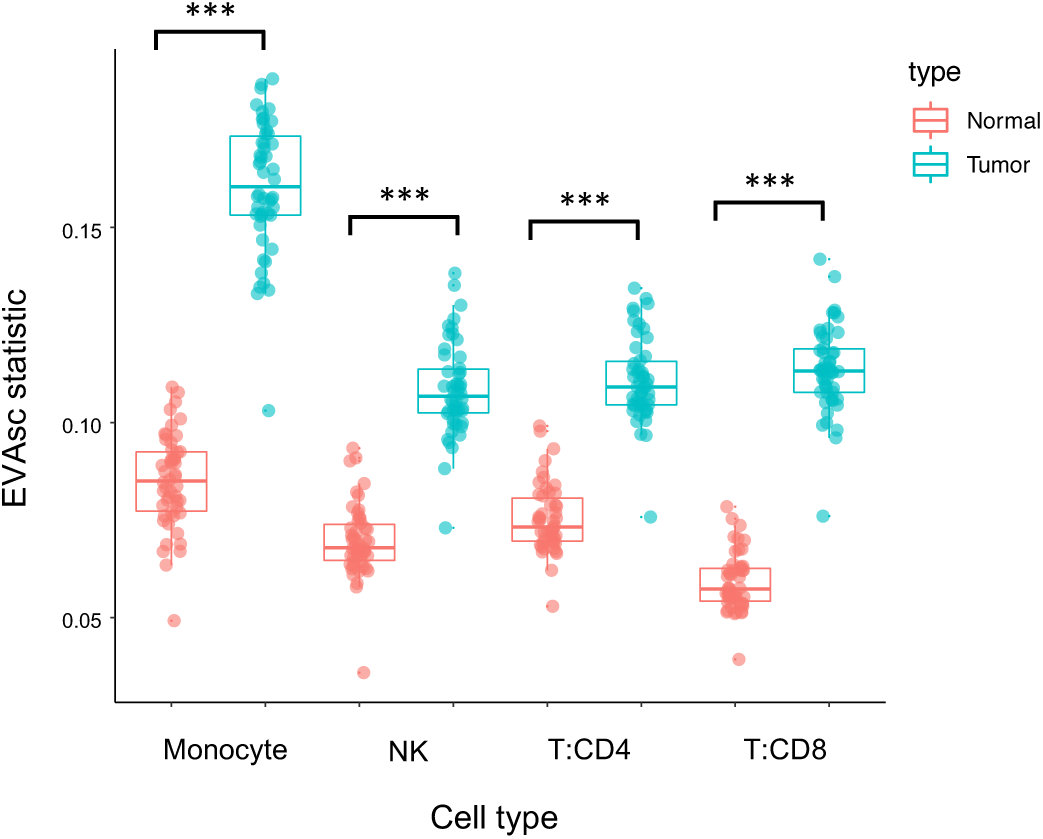
All pathways are significantly dysregulated in immune cell types from breast tumors relative to normal breast tissue. Boxplot of EVAsc statistics of inter-cellular heterogeneity for all fifty hallmark path-ways in major immune cell types from both tumor (blue) and normal (red) breast tissue.

### 3.4 EVAsc finds increased immune pathway heterogeneity in tumors with high T-cell clonality

With the rapid increase of interest in the field of immunotherapy, T-cell receptor (TCR) sequencing is becoming a valuable tool for assessing immune response. Accordingly, we used T-cells from breast cancer data^21^ to explore the relationship between the TCR repertoire and heterogeneity in immune signaling pathways using 27,000 T-cells with paired single-cell RNA and V(D)J sequencing from three breast cancer tumors. For each individual tumor, we computed Shannon entropy for TCR clonality and richness as a measure of TCR diversity based on the single-cell TCR sequencing data (Figure 4A). A Morisita-Horn similarity matrix was generated to compare the similarity of TCR repertoires across tumor replicates (Figure 4B). We then applied EVAsc to the scRNA-seq data using the Myeloid Innate Immunity Panel pathways from NanoString^®^ to compare each T-cell subtype between individuals. Hierarchical clustering of the EVA statistics revealed a gradient of pathway dysregulation directly correlated with the degree of TCR clonality (Figure 4C, Supplemental Table 5). We further applied GSEA to differentially expressed genes to compare the overlap between the enrichment of upregulated and downregulated immune pathways and the immune pathway dysregulation found with EVAsc (Supplemental Tables 6-7). The majority of the significantly dysregulated pathways from EVAsc overlapped with pathways that were enriched for upregulation in higher clonality compared to lower clonality individuals, with seven additional pathway comparisons that are significantly downregulated. We note that clonal expansion of T-cells is generally associated with a mounting immune response after antigen recognition. Our EVAsc results suggest that increased clonality of the TCR repertoire leads to increased heterogeneity in immune pathway expression as well as upregulated immune pathway expression.

**Figure 4.**
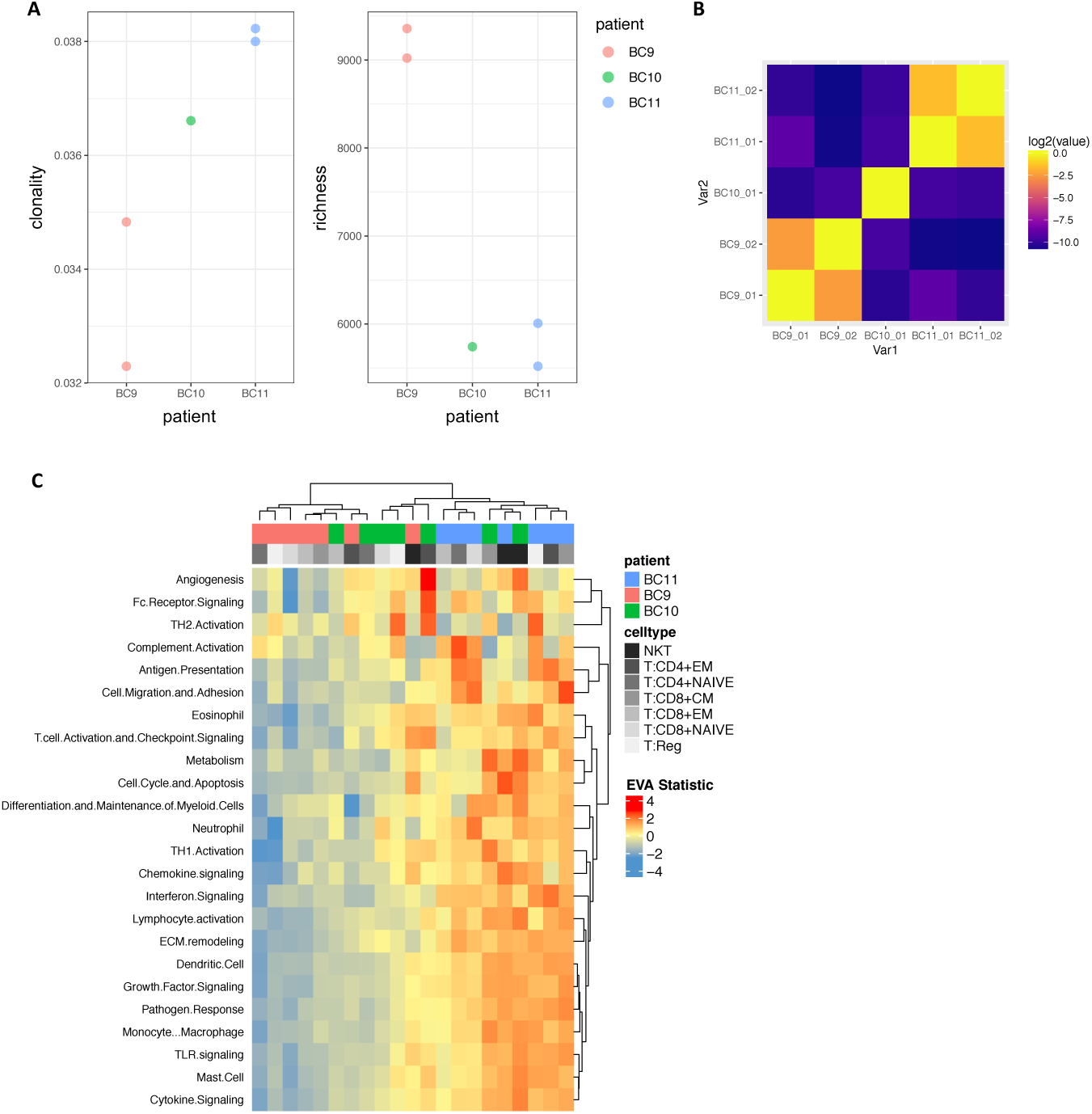
TCR clonality is associated with immune pathway dysregulation in breast tumors. A. TCR clonality and richness for individual breast tumors with matched scRNA-seq and TCR-seq data. B. Heatmap of Morisita-Horn similarity index to quantify agreement of CDR3 clonotypes from duplicate TCR-seq data for the same breast tumor and between individual breast tumors. C. Hierarchical heatmap of EVAsc statistics of inter-cellular heterogeneity for immune pathways in each breast tumor T-cell subtype.

### 3.5 EVAsc finds increased variation in primary tumors relative to metastases and sub-type-specific pathway dysregulation

After demonstrating the ability of EVAsc to detect heterogeneity between tumors, we next aimed to identify differences in pathway heterogeneity between cancer subtypes, within subtypes, and within the TME. Further, we sought to characterize intra-tumor heterogeneity within primary tumors and associated metastases. In order to make these comparisons, we applied EVAsc to scRNA-seq data for 18 HNSCC patients, including five matched primary tumors and lymph node metastases^23^.

We first applied EVAsc to primary cancer cells of HNSCC subtypes to examine the differences between inter-tumor heterogeneity. Subtypes were previously called by TCGA classification and ComBat^33^ was performed to remove the impact of patient identity on transcriptional profiles to compare cells from several patients with EVAsc^29^. We include all MSigDB Hallmark gene set pathways and six meta-signatures derived from non-negative matrix factorization programs that represent common expression programs variable within multiple tumor forms^23^ in our comparisons. 46% (77 out of 168) of the comparisons are statistically significant when all pairwise combinations of subtypes were considered (FDR adjusted p-value < 0.05) (Supplemental Table 8). Hierarchical clustering of the EVAsc statistics demonstrated patterns of subtype-specific pathway dysregulation (Figure 5A).

**Figure 5.**
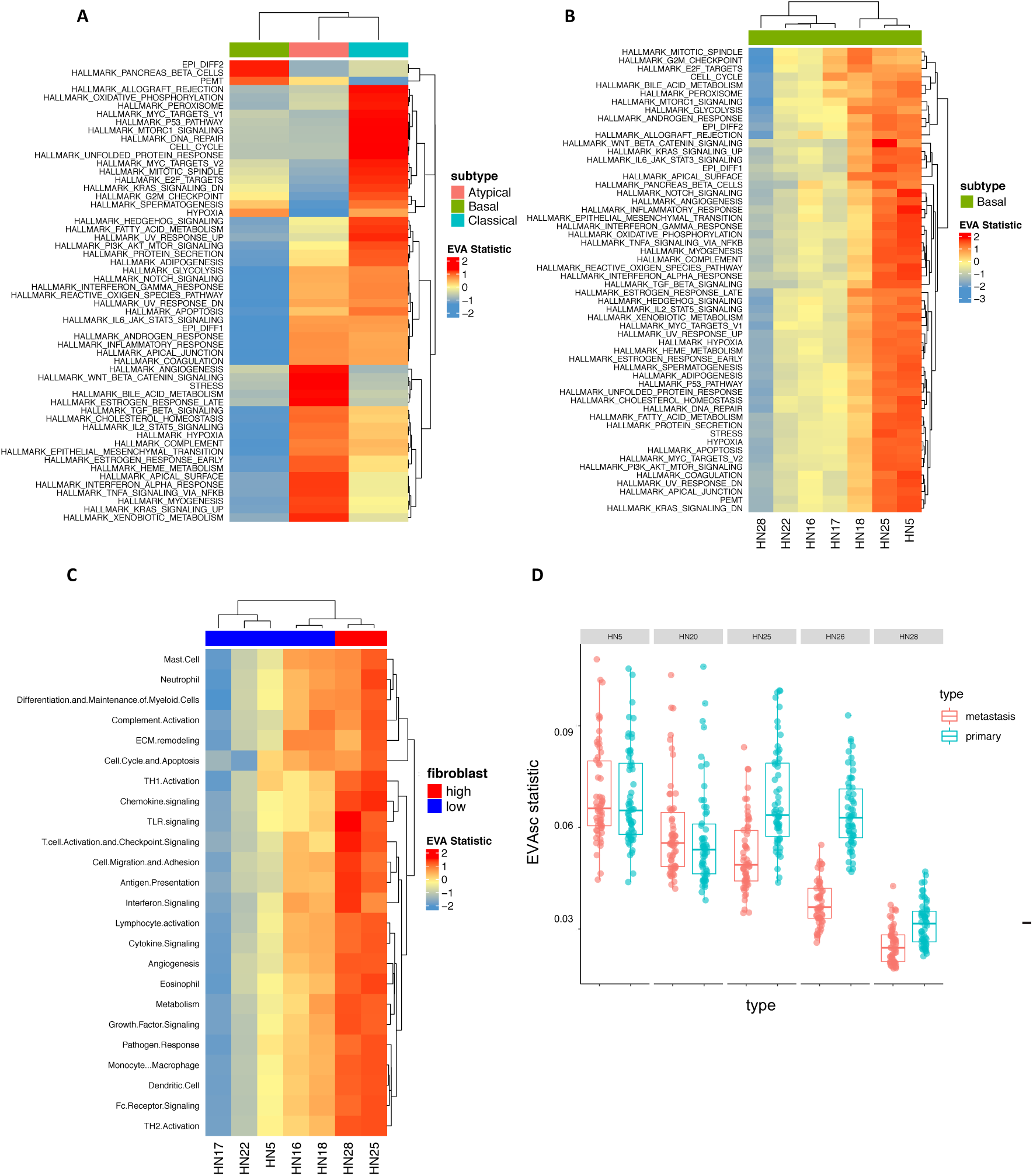
Inter-and intra-tumor heterogeneity distinguish HNSCC subtypes and metastases. A. Heatmap of EVAsc statistics of inter-cellular heterogeneity in hallmark pathways for cancer cells from patients in distinct HNSCC subtypes. B. A heatmap of EVAsc statistics reveals that in-ter-cellular heterogeneity varies between primary cancer cells of the basal tumor type for all hallmark pathways, although no differences in mean expression were observed previously with tSNE^23^. C. EVAsc analysis observes significant increases in inter-cellular variation of immune pathways for fibroblasts that are associated with the total fibroblast content in each basal HNSCC tumor. Previous observations of TCGA sub-types noted that tumors with high fibroblast content (red) were classified as mesenchymal and low fibroblast content (blue) as basal, suggestive of fibroblast mediated differences between immune pathway activity in these subtypes. D. Box-plot of EVAsc statistics in primary and metastatic HNSCC cancer cells for each patient demonstrate higher inter-cellular heterogeneity in primary cancer cells than metastatic cells for two patients.

To determine the degree of inter-tumor heterogeneity between patients within a single subtype we compared primary cancer cells between seven individuals with basal primary tumors. 78% (923 out of 1176) of the comparisons are statistically significant when all pairwise combinations of patients were considered (FDR adjusted p-value < 0.05) (Supplemental Table 9). EVAsc analysis revealed dramatic differences in pathway dysregulation across patients (Figure 5B). Additionally, we explored heterogeneity within cells of the primary TME across individuals with basal primary tumors. Previously, Puram et al.^23^ observed that the proportion of cell types within the TME vary for each patient. Notably, they found that the differences in the basal and mesenchymal subtypes of HPV-negative head and neck cancer can be attributed to a larger proportion of fibroblasts in the TME. Thus, we stratified these basal samples into a binary classification of high (>40%) or low-fibroblast (<40%). To determine the transcriptional status of immune-pathways within patient-specific populations of fibroblasts we applied EVAsc using the Myeloid Innate Immunity Panel pathways from NanoString^®^. Hierarchical clustering of the EVAsc statistics demonstrated increased immune dysregulation in individuals with a high proportion of fibroblasts present in the TME (Figure 5C). 69% (348 out of 504) of the comparisons are statistically significant when all pairwise combinations of patients were considered (FDR adjusted p-value < 0.05) (Supplemental Table 10). We note that the fibroblast composition in each basal tumor is independent of the pathway dysregulation observed across cancer cells from distinct patients.

We next examined intra-tumor pathway dysregulation between the matched primary and metastatic cancer cells within five individual HNSCC patients. 98% (55 out of 56) of the pathways are statistically significant for patient HN25, with 100% (56 out of 56) statistically significant for patient HN26 (FDR adjusted p-value < 0.05) (Supplemental Table 11). In both cases, all significant hits have greater variation in the primary tumor than the metastasis (Figure 5D). For the remaining three patients, no significant pathway dysregulation was observed. Puram et al.^23^ previously observed that the expression profiles of lymph node metastases overlapped with the corresponding primary tumors. While this indicates that there appears to be no mean differences between the paired samples, our method is able to capture significant differential variation between these phenotypes which was previously unrecognized.

## 4. DISCUSSION

We develop EVAsc to quantify heterogeneity in pathway level gene expression from imputed scRNA-seq data to quantify differential variability between conditions. We demonstrate the suitability of EVAsc for identifying differential variability of pathway gene expression by applying it to simulated and real scRNA-seq data. Simulated data generated with splatter was used to demonstrate the ability EVAsc to detect known variability between conditions. Validation was performed by comparing immune cell types between normal breast tissue and breast tumors from Azizi et al.^21^. As expected, EVAsc detected increased variability in the tumor cells for all cell type comparisons relative to normal cells (Figure 3). We then applied EVAsc to perform novel analyses of differential heterogeneity on two publicly available cancer scRNA-seq datasets. We used paired single-cell RNA and single-cell TCR sequencing data^21^ to compare inter-patient T-cell subtype heterogeneity in relation to TCR clonality. TCR repertoire analysis showed differences in the level of TCR clonality for each individual (Figure 4A). EVAsc analysis revealed significant differences in immune pathway heterogeneity between individuals, consistent with the degree of TCR clonality: increased TCR clonality, increased heterogeneity (Figure 4C). We then performed differential expression analysis between individuals to explore the direction of gene set enrichment. There was a large amount of overlap in differentially variable and differentially upregulated pathways, indicating increased heterogeneity as well as increased gene expression in higher clonality individuals.

Ikeda et al.^34^ examined the relationship between intra-tumor expression levels of immune-related genes and TCR repertoire in endometrial cancer. They found increased mRNA expression levels in cases with high T-cell clonality, which was associated with a better prognosis. These results were obtained using total RNA and Quantitative real-time PCR in relatively few genes and are consistent with our findings at a comprehensive single-cell RNA level. Recent data has also shown that increased clonal expansion of T-cells and low baseline clonality are associated with longer survival after being treated with anti-CTLA4 inhibitors in pancreatic ductal adenocarcinoma^30^. Thus, characterizing the immune microenvironment by expression of immune pathways, immune pathway heterogeneity, and the clonality of infiltrated T-cell receptors may be an important biomarker for clinical response to immunotherapy. With the advent of paired single-cell RNA and TCR profiling methods, studying the transcriptional effect of TCR repertoire changes across cancer cells may provide further insight into the mechanisms of immunotherapy.

Further, an HNSCC cancer scRNA-seq dataset from Puram et al.^23^ was used to examine differences in heterogeneity between HNSCC subtypes. Previously, bulk studies have classified HNSCC tumors into four distinct molecular subtypes based on their expression profiles^28^: atypical, basal, classical, and mesenchymal. EVAsc analysis revealed unique patterns of pathway dysregulation in each of the subtypes detected by TCGA classification (Figure 5A). Overall, immune pathways are enriched in the atypical subtype. It has been reported that mesenchymal and atypical subtypes have the highest degree of immune infiltration, making them attractive targets for immunotherapy^35^. Our results suggest a key immune component specific to the atypical subtype.

Previous analyses of the HNSCC scRNA-seq data found that the cancer cells in the mesenchymal and basal subtypes have similar expression profiles when stromal contribution was removed^23^ and refined the classification of mesenchymal to basal subtype. We speculated that the cellular compositions of the TME within individual basal tumors could contribute to the molecular heterogeneity. Importantly, fibroblasts have opposing roles in the TME and showed a wide-range of inter-tumor proportional variability. Normal fibroblasts exert anti-tumorigenic effects to suppress tumor growth but can be reprogrammed to a cancer-associated phenotype supportive of tumor evolution. EVAsc analysis comparing fibroblast populations between individuals with basal primary tumors demonstrated that TMEs with a large proportion of fibroblasts have a high degree of immune pathway dysregulation. This indicated immune pathway heterogeneity within the fibroblast expression states, likely due to the immunomodulatory role of cancer-associated fibroblasts within the TME^36^.

Beyond immunology, the intra-patient comparison with EVAsc enables evaluation of the role of intra-tumor heterogeneity in metastasis. Specifically, we compared cancer cells from primary tumors to metastases from individual patients in HNSCC single-cell data. This analysis revealed a clear pattern: either uniform dysregulation or no significant differences between the primary tumor and metastasis. For the two patients that had differential variability, the heterogeneity within the primary tumor was significantly higher than the metastatic cancer cells (Figure 5D). This observation agrees with Nowell’s theory of clonal evolution, which states that cancer originates from a single cell, accumulates genetic alterations, and during the process of metastasis there is an enrichment for the most aggressive clones^37^. This theory would indicate that clonal metastases are more homogeneous, as very few cells gain invasive and metastatic potential. Such intra-tumor discrepancies that may evolve as the disease progresses between the primary tumor and disseminated metastasis can result in incorrect biomarkers being used to make clinical decisions and lead to therapeutic failure^1^. The differences in molecular heterogeneity may also give rise to different therapeutic responses in primary tumors than metastases. We note that the analyses performed in this study used current landmark cohorts of breast and head and neck tumors, which were limited in sample size. Future work with EVA analysis on larger sample cohorts is essential to establish the role of heterogeneity in complex dynamic processes in cancer progression and therapeutic response.

Together, the results of these analyses show that EVAsc is a robust algorithm for detecting inter-and intra-tumor heterogeneity in scRNA-seq data. EVAsc is applicable to imputed scRNA-seq datasets, which we demonstrate using MAGIC and BiSCUIT imputed data. In the applications to some of the cancer datasets in this study, such as tumor versus normal and primary tumor versus metastasis, we observe widespread changes across a majority of pathways between phenotypes. We attribute these changes to the pervasive transcriptional reprogramming in cancer. While the pathways examined in this study are in no way exhaustive, this is suggestive of global disruption of gene expression and makes for broad interpretations. Comparisons within immune cells and primary cancer subtypes show phenotype specific patterns of dysregulation, allowing more specific interpretation of the molecular mechanisms in tumor heterogeneity. We note that EVAsc can be widely applied beyond cancer, for example to evaluate the role of transcriptional variation on cell fate specification in development^38^. In this context, heterogeneity is more constrained than in cancer and EVAsc finds different patterns of inter-cellular heterogeneity for distinct pathways, with some pathways increasing over developmental time and others decreasing.

In addition, EVAsc is broadly applicable to any dissimilarity metric and is not limited to Kendall-tau (Supplemental Figure 1). This flexibility in the algorithm allows users to specify appropriate distance metrics for datasets and enables the direct comparison of performance across various metrics. We note that different dissimilarity metrics may have different sensitivities to the preprocessing used for the scRNA-seq datasets. The rank-based Kendall-tau dissimilarity metric used for the majority of this study is independent of many sample-specific normalization procedures, such as log transformation or quantile normalization. Other dissimilarity measures may be sensitive to these transformations, and this effect must be evaluated before applying EVAsc to compare dissimilarity based upon these metrics. Emerging variance stabilization methods to account for the pervasive heteroscedastic mean variance relationship of scRNA-seq data may impact the results obtained with this algorithm, and are essential to evaluate in future studies. Thus, EVAsc is a robust multivariate statistical method to quantify differential variation of pathway gene expression and provides the ability to explore transcriptional variation in numerous disease and normal contexts at a single-cell resolution. Future work to improve the EVAsc algorithm will involve integrating mathematical models to compute comparisons on scRNA-seq data without the need for imputation.

## Supporting information

## ACKNOWLEDGEMENTS

This work was supported by grants from the NIH (R01CA177669, U01CA196390, and U01CA212007 to EJF), the Chan-Zuckerberg Initiative DAF (2018-183445 to LAG and 2018-183444 to EJF) an advised fund of Silicon Valley Community Foundation, the Johns Hopkins University Catalyst (EF & LAG) and Discovery awards (EJF), and the Johns Hopkins University School of Medicine Synergy Award (LAG, & EJF). EMJ and ERT acknowledge funding from the Broccoli Foundation, The Bloomberg~Kimmel Institute for Cancer Immunotherapy, and The Skip Viragh Center for Pancreas Cancer Clinical Research and Patient Care, and The Commonwealth Foundation for Cancer Research. ERT is funded through the MacMillan Pathway to Independence Fellowship. EMJ, ERT, and AH are also supported through NIH R01CA184926, as well as Stand Up To Cancer which is a program of the Entertainment Industry Foundation administered by the American Association for Cancer Research. The authors thank L. Cope, A. Ewald, K. Schuebel, R. Scharpf, V. Yegnasu-bramanian, R. Riggins, L. Kagohara, D. Gaykalova, T. Triche, and W. H. Jin for feedback on the algorithm and manuscript.

---

**Supplemental Figure 1.**
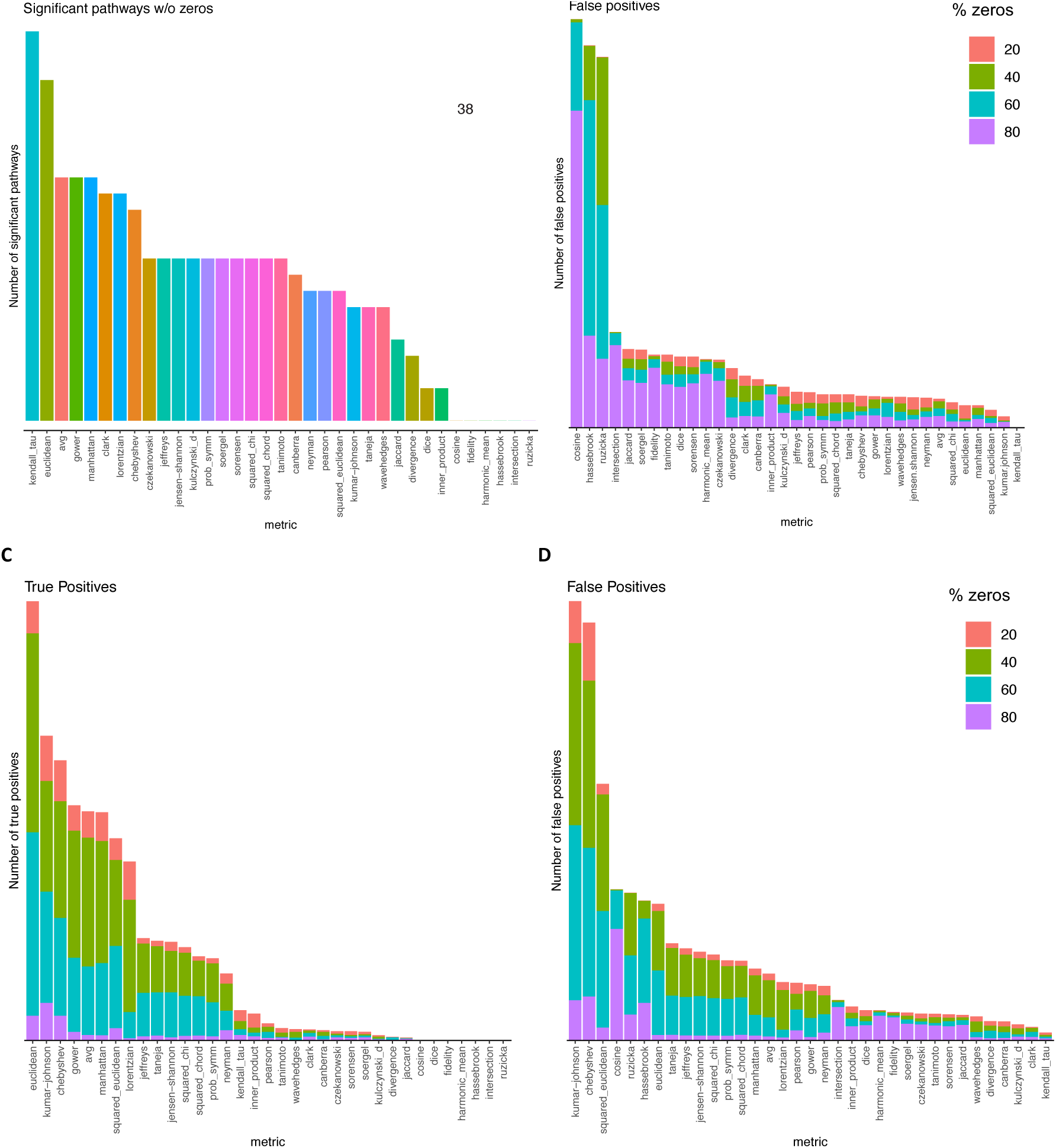
A. Barplot of the number of significant hallmark gene set pathways across 35 metrics from performing EVA on a benchmark bulk RNA-sequencing dataset with normal and cancer samples. B. Number of significant hallmark gene set pathways across 35 metrics from performing 100 EVA permutations on a mirrored bulk RNA-sequencing data set so that there is no signal. Each bar is colored by the percentage of random zeros added to the dataset. As the number of random zeros increases, the number of significant pathways (all false positives) tends to increase. C. Number of true significant hallmark gene set pathways across 35 metrics from performing 100 EVA permutations on the bulk RNA-sequencing data from (A). True positives are defined as any pathway that was significant for a metric when no zeros are present. Each bar is colored by the percentage of random zeros added to the dataset. For most metrics, as the number of random zeros increases, the number of significant pathways increases around 40% and 60% zeros, and the signal drops at 80%. D> Number of false positive significant hallmark gene set pathways across 35 metrics from performing 100 EVA permutations on the bulk RNA-sequencing data from (A). False positives are defined as any pathway that was not significant for a metric when no zeros are present. Each bar is colored by the percentage of random zeros added to the dataset. For most metrics, as the number of random zeros increases, the number of significant pathways peaks around 40% and 60% zeros, with some metrics peaking at 80%. This indicates variable sensitivity to zeros in a dataset with signal.

