## Supplementary material for "Expression variation analysis for tumor heterogeneity in single-cell RNA-sequencing data"

| | gst | pathway_name | sample1 | sample2 | celltype | gsea |
|---|---|---|---|---|---|---|
| 31 | 2.507837e−02 | Cytokine.Signaling | BC10 | BC11 | T:CD4+EM | up |
| 35 | 3.246457e−03 | Growth.Factor.Signaling | BC10 | BC11 | T:CD4+EM | up |
| 36 | 7.333910e−06 | Interferon.Signaling | BC10 | BC11 | T:CD4+EM | up |
| 37 | 1.504184e−03 | Lymphocyte.activation | BC10 | BC11 | T:CD4+EM | up |
| 41 | 7.156757e−03 | TH1.Activation | BC10 | BC11 | T:CD4+EM | up |
| 42 | 3.448037e−02 | TH2.Activation | BC10 | BC11 | T:CD4+EM | up |
| 47 | 4.441081e−05 | Dendritic.Cell | BC10 | BC11 | T:CD4+EM | up |
| 48 | 7.815302e−03 | Monocyte...Macrophage | BC10 | BC11 | T:CD4+EM | up |
| 50 | 3.848638e−04 | Antigen.Presentation | BC9 | BC11 | T:CD4+EM | up |
| 55 | 4.947781e−02 | Cytokine.Signaling | BC9 | BC11 | T:CD4+EM | up |
| 57 | 1.342585e−02 | ECM.remodeling | BC9 | BC11 | T:CD4+EM | up |
| 59 | 1.220989e−03 | Growth.Factor.Signaling | BC9 | BC11 | T:CD4+EM | up |
| 60 | 1.328724e−05 | Interferon.Signaling | BC9 | BC11 | T:CD4+EM | up |
| 65 | 5.082992e−03 | TH1.Activation | BC9 | BC11 | T:CD4+EM | up |
| 70 | 1.936704e−02 | Mast.Cell | BC9 | BC11 | T:CD4+EM | up |
| 71 | 7.164126e−04 | Dendritic.Cell | BC9 | BC11 | T:CD4+EM | up |
| 72 | 1.630868e−05 | Monocyte...Macrophage | BC9 | BC11 | T:CD4+EM | up |
| 77 | 1.308780e−02 | Chemokine.signaling | BC9 | BC10 | T:CD4+NAIVE | up |
| 81 | 1.174392e−03 | ECM.remodeling | BC9 | BC10 | T:CD4+NAIVE | up |
| 89 | 5.371754e−03 | TH1.Activation | BC9 | BC10 | T:CD4+NAIVE | up |
| 92 | 4.659071e−02 | Neutrophil | BC9 | BC10 | T:CD4+NAIVE | up |
| 93 | 4.113501e−02 | Eosinophil | BC9 | BC10 | T:CD4+NAIVE | up |
| 96 | 7.593873e−03 | Monocyte...Macrophage | BC9 | BC10 | T:CD4+NAIVE | up |
| 98 | 3.353797e−03 | Antigen.Presentation | BC10 | BC11 | T:CD4+NAIVE | up |
| 101 | 2.762426e−02 | Chemokine.signaling | BC10 | BC11 | T:CD4+NAIVE | up |
| 103 | 5.398627e−03 | Cytokine.Signaling | BC10 | BC11 | T:CD4+NAIVE | up |
| 106 | 4.810486e−02 | Fc.Receptor.Signaling | BC10 | BC11 | T:CD4+NAIVE | up |
| 107 | 2.064575e−03 | Growth.Factor.Signaling | BC10 | BC11 | T:CD4+NAIVE | up |
| 108 | 2.100643e−05 | Interferon.Signaling | BC10 | BC11 | T:CD4+NAIVE | up |
| 109 | 2.185136e−02 | Lymphocyte.activation | BC10 | BC11 | T:CD4+NAIVE | up |
| 113 | 1.342226e−03 | TH1.Activation | BC10 | BC11 | T:CD4+NAIVE | up |
| 114 | 4.597082e−03 | TH2.Activation | BC10 | BC11 | T:CD4+NAIVE | up |
| 116 | 3.762637e−04 | Neutrophil | BC10 | BC11 | T:CD4+NAIVE | up |
| 118 | 1.180163e−02 | Mast.Cell | BC10 | BC11 | T:CD4+NAIVE | up |
| 119 | 4.160929e−06 | Dendritic.Cell | BC10 | BC11 | T:CD4+NAIVE | up |
| 120 | 1.563977e−05 | Monocyte...Macrophage | BC10 | BC11 | T:CD4+NAIVE | up |
| 122 | 1.019635e−03 | Antigen.Presentation | BC9 | BC11 | T:CD4+NAIVE | up |
| 125 | 2.375596e−03 | Chemokine.signaling | BC9 | BC11 | T:CD4+NAIVE | up |
| 127 | 2.816396e−04 | Cytokine.Signaling | BC9 | BC11 | T:CD4+NAIVE | up |
| 129 | 8.247648e−04 | ECM.remodeling | BC9 | BC11 | T:CD4+NAIVE | up |
| 131 | 7.185803e−05 | Growth.Factor.Signaling | BC9 | BC11 | T:CD4+NAIVE | up |
| 132 | 8.890153e−06 | Interferon.Signaling | BC9 | BC11 | T:CD4+NAIVE | up |
| 134 | 3.375217e−03 | Metabolism | BC9 | BC11 | T:CD4+NAIVE | up |
| 135 | 2.920299e−02 | Pathogen.Response | BC9 | BC11 | T:CD4+NAIVE | up |
| 137 | 4.440731e−05 | TH1.Activation | BC9 | BC11 | T:CD4+NAIVE | up |
| 139 | 3.374153e−02 | TLR.signaling | BC9 | BC11 | T:CD4+NAIVE | up |
| 140 | 6.289460e−04 | Neutrophil | BC9 | BC11 | T:CD4+NAIVE | up |
| 141 | 2.745515e−04 | Eosinophil | BC9 | BC11 | T:CD4+NAIVE | up |
| 142 | 9.431077e−04 | Mast.Cell | BC9 | BC11 | T:CD4+NAIVE | up |
| 143 | 4.357071e−09 | Dendritic.Cell | BC9 | BC11 | T:CD4+NAIVE | up |
| 144 | 2.360092e−09 | Monocyte...Macrophage | BC9 | BC11 | T:CD4+NAIVE | up |
| 148 | 3.743150e−02 | Cell.Migration.and.Adhesion | BC9 | BC10 | T:CD8+CM | up |
| 151 | 9.677036e−03 | Cytokine.Signaling | BC9 | BC10 | T:CD8+CM | up |
| 155 | 2.590994e−02 | Growth.Factor.Signaling | BC9 | BC10 | T:CD8+CM | up |
| 156 | 2.033433e−02 | Interferon.Signaling | BC9 | BC10 | T:CD8+CM | up |
| 168 | 2.267636e−02 | Monocyte...Macrophage | BC9 | BC10 | T:CD8+CM | up |
| 180 | 2.152378e−03 | Interferon.Signaling | BC10 | BC11 | T:CD8+CM | up |
| 185 | 3.771985e−02 | TH1.Activation | BC10 | BC11 | T:CD8+CM | up |
| 186 | 7.557016e−03 | TH2.Activation | BC10 | BC11 | T:CD8+CM | up |
| 191 | 4.372187e−02 | Dendritic.Cell | BC10 | BC11 | T:CD8+CM | up |
| 199 | 1.907095e−02 | Cytokine.Signaling | BC9 | BC11 | T:CD8+CM | up |
| 203 | 2.219588e−02 | Growth.Factor.Signaling | BC9 | BC11 | T:CD8+CM | up |
| 204 | 1.232415e−03 | Interferon.Signaling | BC9 | BC11 | T:CD8+CM | up |
| 209 | 2.405159e−02 | TH1.Activation | BC9 | BC11 | T:CD8+CM | up |
| 215 | 4.369093e−03 | Dendritic.Cell | BC9 | BC11 | T:CD8+CM | up |
| 216 | 8.900209e−03 | Monocyte...Macrophage | BC9 | BC11 | T:CD8+CM | up |
| 219 | 4.541629e−02 | Cell.Cycle.and.Apoptosis | BC9 | BC10 | T:CD8+EM | up |
| 223 | 2.119113e−03 | Cytokine.Signaling | BC9 | BC10 | T:CD8+EM | up |
| 227 | 6.646961e−04 | Growth.Factor.Signaling | BC9 | BC10 | T:CD8+EM | up |
| 229 | 1.260704e−02 | Lymphocyte.activation | BC9 | BC10 | T:CD8+EM | up |
| 238 | 6.177730e−03 | Mast.Cell | BC9 | BC10 | T:CD8+EM | up |
| 242 | 6.483536e−04 | Antigen.Presentation | BC10 | BC11 | T:CD8+EM | up |
| 251 | 1.825078e−02 | Growth.Factor.Signaling | BC10 | BC11 | T:CD8+EM | up |
| 252 | 4.307121e−04 | Interferon.Signaling | BC10 | BC11 | T:CD8+EM | up |
| 254 | 3.358726e−02 | Metabolism | BC10 | BC11 | T:CD8+EM | up |
| 255 | 1.166250e−03 | Pathogen.Response | BC10 | BC11 | T:CD8+EM | up |
| 257 | 1.009401e−02 | TH1.Activation | BC10 | BC11 | T:CD8+EM | up |
| 258 | 4.749872e−02 | TH2.Activation | BC10 | BC11 | T:CD8+EM | up |
| 262 | 1.330620e−02 | Mast.Cell | BC10 | BC11 | T:CD8+EM | up |
| 263 | 5.339219e−04 | Dendritic.Cell | BC10 | BC11 | T:CD8+EM | up |
| 264 | 1.442057e−04 | Monocyte...Macrophage | BC10 | BC11 | T:CD8+EM | up |
| 266 | 1.856957e−03 | Antigen.Presentation | BC9 | BC11 | T:CD8+EM | up |
| 267 | 7.085716e−03 | Cell.Cycle.and.Apoptosis | BC9 | BC11 | T:CD8+EM | up |
| 271 | 7.964382e−05 | Cytokine.Signaling | BC9 | BC11 | T:CD8+EM | up |
| 275 | 6.434673e−06 | Growth.Factor.Signaling | BC9 | BC11 | T:CD8+EM | up |
| 276 | 1.023317e−05 | Interferon.Signaling | BC9 | BC11 | T:CD8+EM | up |
| 279 | 6.016659e−03 | Pathogen.Response | BC9 | BC11 | T:CD8+EM | up |
| 281 | 1.008764e−03 | TH1.Activation | BC9 | BC11 | T:CD8+EM | up |
| 284 | 1.174729e−03 | Neutrophil | BC9 | BC11 | T:CD8+EM | up |
| 285 | 1.196129e−02 | Eosinophil | BC9 | BC11 | T:CD8+EM | up |
| 286 | 1.404797e−02 | Mast.Cell | BC9 | BC11 | T:CD8+EM | up |
| 287 | 4.841109e−05 | Dendritic.Cell | BC9 | BC11 | T:CD8+EM | up |
| 288 | 8.182150e−07 | Monocyte...Macrophage | BC9 | BC11 | T:CD8+EM | up |
| 297 | 1.667742e−02 | ECM.remodeling | BC9 | BC10 | T:CD8+NAIVE | up |
| 305 | 3.692216e−02 | TH1.Activation | BC9 | BC10 | T:CD8+NAIVE | up |
| 314 | 1.185721e−02 | Antigen.Presentation | BC10 | BC11 | T:CD8+NAIVE | up |
| 319 | 5.480575e−03 | Cytokine.Signaling | BC10 | BC11 | T:CD8+NAIVE | up |
| 322 | 4.159795e−02 | Fc.Receptor.Signaling | BC10 | BC11 | T:CD8+NAIVE | up |
| 323 | 1.358619e−03 | Growth.Factor.Signaling | BC10 | BC11 | T:CD8+NAIVE | up |
| 324 | 7.538080e−05 | Interferon.Signaling | BC10 | BC11 | T:CD8+NAIVE | up |
| 330 | 1.017888e−02 | TH2.Activation | BC10 | BC11 | T:CD8+NAIVE | up |
| 334 | 3.273970e−03 | Mast.Cell | BC10 | BC11 | T:CD8+NAIVE | up |
| 335 | 4.796281e−04 | Dendritic.Cell | BC10 | BC11 | T:CD8+NAIVE | up |
| 336 | 6.577703e−03 | Monocyte...Macrophage | BC10 | BC11 | T:CD8+NAIVE | up |
| 343 | 1.498735e−04 | Cytokine.Signaling | BC9 | BC11 | T:CD8+NAIVE | up |
| 347 | 3.061251e−04 | Growth.Factor.Signaling | BC9 | BC11 | T:CD8+NAIVE | up |
| 348 | 1.007377e−04 | Interferon.Signaling | BC9 | BC11 | T:CD8+NAIVE | up |
| 351 | 2.074866e−02 | Pathogen.Response | BC9 | BC11 | T:CD8+NAIVE | up |
| 353 | 1.214534e−03 | TH1.Activation | BC9 | BC11 | T:CD8+NAIVE | up |
| 357 | 2.542070e−03 | Eosinophil | BC9 | BC11 | T:CD8+NAIVE | up |
| 358 | 1.118333e−02 | Mast.Cell | BC9 | BC11 | T:CD8+NAIVE | up |
| 359 | 5.518259e−06 | Dendritic.Cell | BC9 | BC11 | T:CD8+NAIVE | up |
| 360 | 1.549707e−04 | Monocyte...Macrophage | BC9 | BC11 | T:CD8+NAIVE | up |
| 363 | 4.683629e−02 | Cell.Cycle.and.Apoptosis | BC9 | BC10 | T:Reg | up |
| 365 | 5.245452e−04 | Chemokine.signaling | BC9 | BC10 | T:Reg | up |
| 367 | 9.789904e−03 | Cytokine.Signaling | BC9 | BC10 | T:Reg | up |
| 371 | 2.895952e−02 | Growth.Factor.Signaling | BC9 | BC10 | T:Reg | up |
| 372 | 2.389114e−02 | Interferon.Signaling | BC9 | BC10 | T:Reg | up |
| 376 | 2.815269e−02 | T.cell.Activation.and.Checkpoint.Signaling | BC9 | BC10 | T:Reg | up |
| 377 | 2.457382e−04 | TH1.Activation | BC9 | BC10 | T:Reg | up |
| 380 | 1.554844e−02 | Neutrophil | BC9 | BC10 | T:Reg | up |
| 382 | 1.902145e−02 | Mast.Cell | BC9 | BC10 | T:Reg | up |
| 383 | 2.105489e−02 | Dendritic.Cell | BC9 | BC10 | T:Reg | up |
| 384 | 1.747533e−02 | Monocyte...Macrophage | BC9 | BC10 | T:Reg | up |
| 386 | 8.398283e−06 | Antigen.Presentation | BC10 | BC11 | T:Reg | up |
| 391 | 1.109369e−03 | Cytokine.Signaling | BC10 | BC11 | T:Reg | up |
| 394 | 3.155234e−02 | Fc.Receptor.Signaling | BC10 | BC11 | T:Reg | up |
| 395 | 1.095006e−03 | Growth.Factor.Signaling | BC10 | BC11 | T:Reg | up |
| 396 | 2.034091e−06 | Interferon.Signaling | BC10 | BC11 | T:Reg | up |
| 397 | 2.091628e−02 | Lymphocyte.activation | BC10 | BC11 | T:Reg | up |
| 402 | 3.186999e−02 | TH2.Activation | BC10 | BC11 | T:Reg | up |
| 405 | 4.350302e−02 | Eosinophil | BC10 | BC11 | T:Reg | up |
| 407 | 6.448286e−04 | Dendritic.Cell | BC10 | BC11 | T:Reg | up |
| 408 | 1.150338e−06 | Monocyte...Macrophage | BC10 | BC11 | T:Reg | up |
| 410 | 6.222312e−05 | Antigen.Presentation | BC9 | BC11 | T:Reg | up |
| 413 | 2.022802e−03 | Chemokine.signaling | BC9 | BC11 | T:Reg | up |
| 415 | 3.210594e−06 | Cytokine.Signaling | BC9 | BC11 | T:Reg | up |
| 419 | 2.283913e−05 | Growth.Factor.Signaling | BC9 | BC11 | T:Reg | up |
| 420 | 1.656737e−06 | Interferon.Signaling | BC9 | BC11 | T:Reg | up |
| 423 | 9.496928e−03 | Pathogen.Response | BC9 | BC11 | T:Reg | up |
| 425 | 5.925661e−04 | TH1.Activation | BC9 | BC11 | T:Reg | up |
| 426 | 1.116948e−02 | TH2.Activation | BC9 | BC11 | T:Reg | up |
| 429 | 1.405182e−04 | Eosinophil | BC9 | BC11 | T:Reg | up |
| 430 | 2.606450e−02 | Mast.Cell | BC9 | BC11 | T:Reg | up |
| 431 | 1.157189e−07 | Dendritic.Cell | BC9 | BC11 | T:Reg | up |
| 432 | 2.014306e−10 | Monocyte...Macrophage | BC9 | BC11 | T:Reg | up |
| 468 | 1.123544e−04 | Interferon.Signaling | BC10 | BC11 | NKT | up |
| 473 | 3.513204e−02 | TH1.Activation | BC10 | BC11 | NKT | up |
| 474 | 2.740002e−02 | TH2.Activation | BC10 | BC11 | NKT | up |
| 479 | 3.407699e−02 | Dendritic.Cell | BC10 | BC11 | NKT | up |
| 491 | 3.127764e−02 | Growth.Factor.Signaling | BC9 | BC11 | NKT | up |
| 492 | 1.317012e−04 | Interferon.Signaling | BC9 | BC11 | NKT | up |
| 493 | 1.604273e−02 | Lymphocyte.activation | BC9 | BC11 | NKT | up |
