## Supplementary material for "Expression variation analysis for tumor heterogeneity in single-cell RNA-sequencing data"

| | gst | pathway_name | sample1 | sample2 | celltype | gsea |
|---|---|---|---|---|---|---|
| *21* | 0.006012080 | Eosinophil | BC9 | BC10 | T:CD4+EM | down |
| *172* | 0.007589262 | Cell.Migration.and.Adhesion | BC10 | BC11 | T:CD8+CM | down |
| *193* | 0.043070623 | Angiogenesis | BC9 | BC11 | T:CD8+CM | down |
| *400* | 0.043505614 | T.cell.Activation.and.Checkpoint.Signaling | BC10 | BC11 | T:Reg | down |
| *437* | 0.010527004 | Chemokine.signaling | BC9 | BC10 | NKT | down |
| *455* | 0.010487495 | Dendritic.Cell | BC9 | BC10 | NKT | down |
| *485* | 0.012289230 | Chemokine.signaling | BC9 | BC11 | NKT | down |
